# Targeted DamID detects cell-type specific histone modifications *in vivo*

**DOI:** 10.1101/2024.04.11.589050

**Authors:** Jelle van den Ameele, Manuel Trauner, Eva Hörmanseder, Alex P. A. Donovan, Oriol Llora Battle, Seth W. Cheetham, Robert Krautz, Rebecca Yakob, John B. Gurdon, Andrea H. Brand

## Abstract

Histone modifications play a key role in regulating gene expression and cell fate during development and disease. Current methods for cell-type specific genome-wide profiling of histone modifications require dissociation and isolation of cells and are not compatible with all tissue types. Here we adapt Targeted DamID to recognise specific histone marks, by fusing chromatin binding proteins or single-chain antibodies to Dam, an *E. coli* DNA adenine methylase. When combined with Targeted DamID (TaDa), this enables cell-type specific chromatin profiling in intact tissues or organisms. We first profiled H3K4me3, H3K9ac, H3K27me3 and H4K20me1 *in vivo* in neural stem cells of the developing *Drosophila* brain. Next, we mapped cell-type specific H3K4me3 distribution in neural stem cells of the developing mouse brain. Finally, we injected RNA encoding DamID constructs into 1-cell stage *Xenopus* embryos to profile H3K4me3 distribution during gastrulation and neurulation. These results illustrate the versatility of Targeted DamID to profile cell-type specific histone marks throughout the genome in diverse model systems.

**Summary statement:** Targeted DamID enables genome-wide cell-type specific detection of histone modifications *in vivo* in *Drosophila*, mouse and *Xenopus*.

## Introduction

Post-translational modifications of histones and chromatin-associated proteins play a crucial role in both normal and pathological cell fate decisions, through their effects on transcription, replication and DNA repair (Bannister and Kouzarides, 2011). There remains a need for tools to profile post-translational chromatin modifications in specific cell types, *in vivo*, without prior cell isolation (Handley et al., 2015). Most current techniques rely on co-immunoprecipitation (ChIP-seq (Johnson et al., 2007)) or tagging (Cut&Run (Skene and Henikoff, 2017)) of DNA associated with the modified histones, and cell sorting is required to obtain cell-type specific information. More recently, ChromID was developed using histone modification specific biotinylated chromatin reader domains fused to GFP, allowing histone modifications to be profiled by ChIP using antibodies against biotin (Villaseñor et al., 2020). Several drawbacks of these approaches include: cross-linking which is biased towards specific regions of chromatin (Landt et al., 2012); cell dissociation, which may cause epigenetic changes (Binek et al., 2018; Potier et al., 2014; van den Brink et al., 2017); immunoprecipitation, which can be difficult in early-stage *Xenopus* embryos as they contain large amounts of cytoplasmic protein as compared to the nucleus.

DamID relies on the recruitment of an *E. coli* DNA adenine methyltransferase (Dam) to specific loci in the genome, where it methylates adenines in nearby GATC-motifs (van Steensel and Henikoff, 2000; Vogel et al., 2007) (**Fig. 1A**). By expressing Dam fused to a chromatin-binding protein of interest, the protein binding sites are marked by G^m6^ATC methylation and can be detected by DamID-seq (Marshall & Brand., 2017). Cell-type specific methylation can be achieved through Targeted DamID (TaDa) (Southall et al., 2013), where the coding sequence of the Dam-fusion protein (ORF2) is inserted downstream of a primary open reading frame (ORF1), thereby reducing the level of Dam expression and avoiding potential toxicity (Southall et al., 2013). Targeted DamID has been used to profile chromatin states, where Dam is fused to proteins that recognise and bind specific histone modifications (Marshall & Brand, 2017). More recently DamID has been used for single cell profiling of a subset of histone marks (H3K27me3, H3K4me3, H4K20me1 and H3K9ac) using either entire chromatin binding proteins, their histone mark binding domains, or histone modification specific mintbodies (single chain variable fragments, scFvs) bound to Dam methylase (EpiDamID, Rang et al., 2022) . However, to assay histone modifications in a cell-type specific manner EpiDamID must be used in combination with single cell sequencing, due to expression from a ubiquitous promoter.

**Figure 1.**
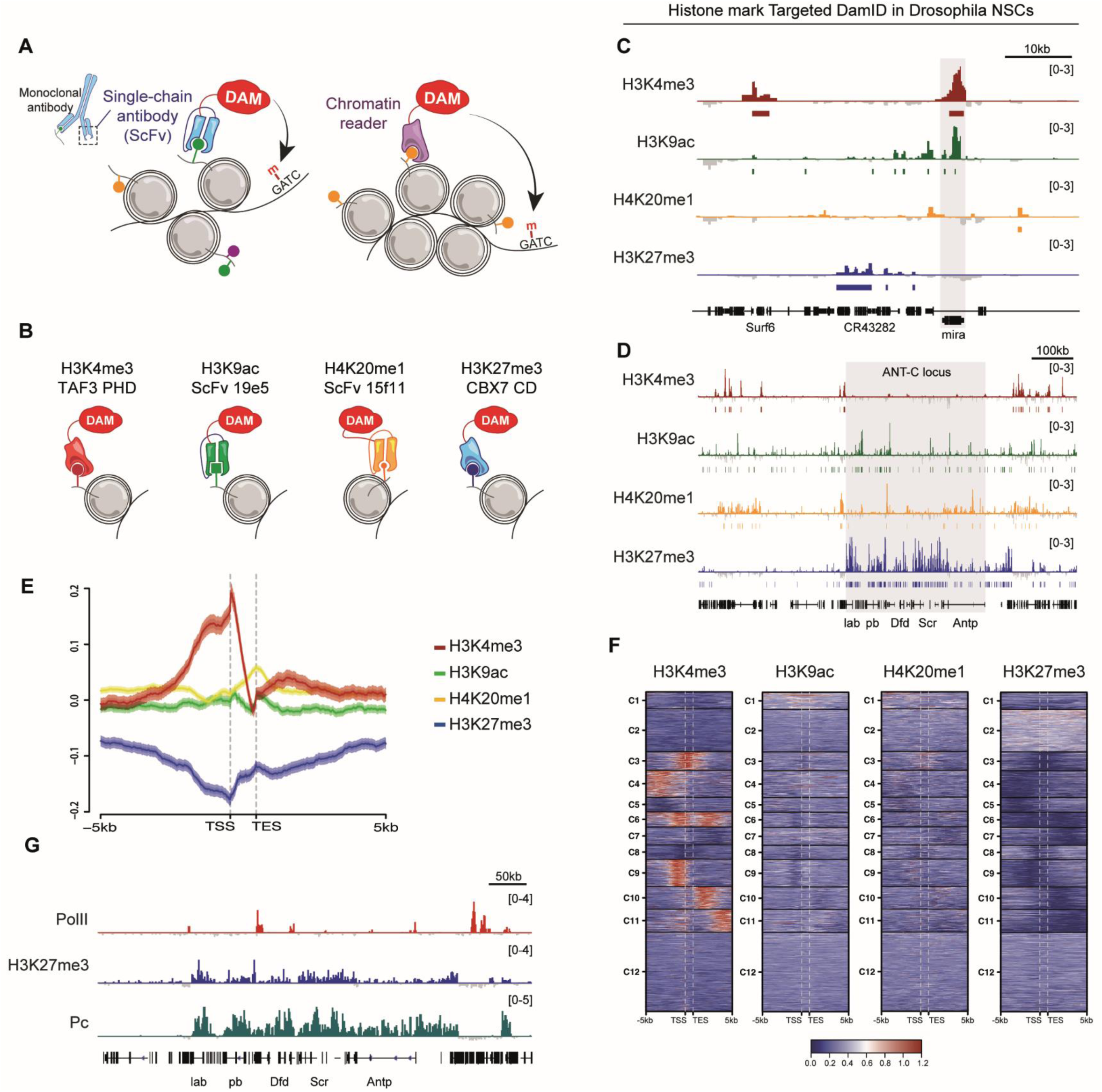
Profiling histone marks with TaDa. (**A**) Schematic showing methods used for Targeted DamID profiling of histone modifications. (**B**) Illustration demonstrating fusion of Dam to chromatin binding domains for H3K4me3 or H3K27me3 and to mintbodies (scFvs) against H3K9ac or H4K20me1. (**C**) Histone marks near the *Mira* locus (shaded) in Drosophila third instar NSCs (**D**) Histone marks near the *ANT-C* locus (shaded) in Drosophila third instar NSCs. (**E**) Average (±s.e.m.) signal intensity and (**F**) intensity of TaDa signal across genes (transcription start/end site, TSS/TES, ±5kb) expressed in third instar larval NSCs. (**G**) TaDa profiles for PolII, H3K27me3 and polycomb (Pc) around the Antennapedia complex.

Here, we describe Targeted DamID for profiling cell-type specific histone marks in intact organisms and demonstrate its efficacy in *Drosophila* neural stem cells (NSCs) in the developing CNS, in NSCs and postmitotic neurons in the developing mouse cerebral cortex and in single *Xenopus* embryos.

## Results

### Profiling histone modifications with TaDa in Drosophila NSCs

We fused Dam to proteins that bind with high affinity and specificity to posttranslational modifications, enabling identification of genome-wide distribution of these marks. We took advantage of several previously characterized mintbodies and protein binding domains (chromatin readers) that recognise histone modifications on native chromatin (**Fig. 1A**). The PHD domain of TAF3 recognises H3K4me3 (Kungulovski et al., 2016), the chromodomain of CBX7 binds H3K27me3 (Kungulovski et al., 2014), the DNMT3A PWWP domain recognises H3K36me3 (Kungulovski et al., 2014) and the single chain monoclonal antibody fragments 19E5 (Sato et al., 2013) and 15E11 (Sato et al., 2016) recognise H3K9ac and H4K20me1 respectively (**Fig. 1B**). All chromatin readers were previously validated by ChIP-seq or peptide arrays (Kungulovski et al., 2016), and the H3K9ac and H4K20me1 mintbodies were optimized to detect histone modifications upon intracellular expression (Mintbodies (Sato et al., 2013; Sato et al., 2016)) and proven compatible with DamID (Rang et al., 2022). We fused these chromatin readers to the *E. coli* Dam methylase, and expressed the Dam-fusions specifically in the NSCs of *Drosophila* third instar larvae using TaDa (Southall et al., 2013). Spatiotemporal control of TaDa expression was achieved with the GAL4 system (Brand and Perrimon, 1993) and GAL80^ts^ (McGuire et al., 2003). All Dam-fusion proteins generated distinct and reproducible methylation profiles (**Fig. 1C,D; Fig. S1A**). As expected, H3K4me3 was enriched in promoter regions, whereas H3K9ac, H4K20me1 and H3K27me3 covered both promoter and coding sequences **(Fig. 1C,D; Fig. S1B)**. Focusing on genes actively transcribed in NSCs at this stage (Marshall & Brand, 2017), H3K4me3 was present at both promoters and coding sequences, while H4K20me1was mainly located in coding sequences. H3K9ac was not prevalent and H3K27me3, which is associated with repressed regions, was absent **(Fig.1E,F)**.

We then compared the NSC-specific binding profiles of the chromatin readers and mintbody with published TaDa of PolII, Brahma, Pc, H1 and HP1a (Marshall & Brand, 2017) to validate the distribution of the profiled histone marks. PolII and H3K4me3 correlated with one another well, as expected, whereas H3K9ac correlated better with Brahma, usually linked to enhancer regions marked by H3K27ac, therefore suggesting that in NSCs H3K9ac might be associated with enhancers rather that the promoters of expressed genes **(Fig.S1C, E)**. In terms of repressive marks, H3K27me3 correlated highly with the Pc and H1 TaDa, indicating that the chromatin reader is correctly recognising the presence of H3K27me3 **(Fig. S1F)**. In fact, this can be clearly observed at loci known to be targeted by Polycomb Group of proteins, such as the Antennapedia complex or vnd locus, where H3K27me3 and Pc binding patterns overlap **(Fig.1G; Fig.S1D)**. Overall, we concluded that our strategy can accurately profile histone marks *in vivo*.

### Profiling histone marks *in vivo* in radial glial cells of the developing mouse cortex

Radial glial cells (RGCs) that line the ventricular zone of the developing mouse cortex can be transiently transfected through injection of plasmid DNA into the telencephalic ventricles of a mouse embryo *in utero*, followed by transcranial electroporation (**Fig. 2A-C**). We fused the chromatin readers for H3K4me3, H3K9ac, H4K20me1 and H3K27me3 to intron-Dam for transient transfection (Wade et al., 2021; van den Ameele et al., 2022) and expressed them downstream of pHes5, an established RGC-specific promoter (Mizutani et al., 2007; Ohtsuka et al., 2006). Histone mark peaks in mouse NSCs were enriched in previously identified by ChIP-seq ( ENCODE Project Consortium., 2012) (**Fig. S2E, F**), validating our approach.

**Figure 2.**
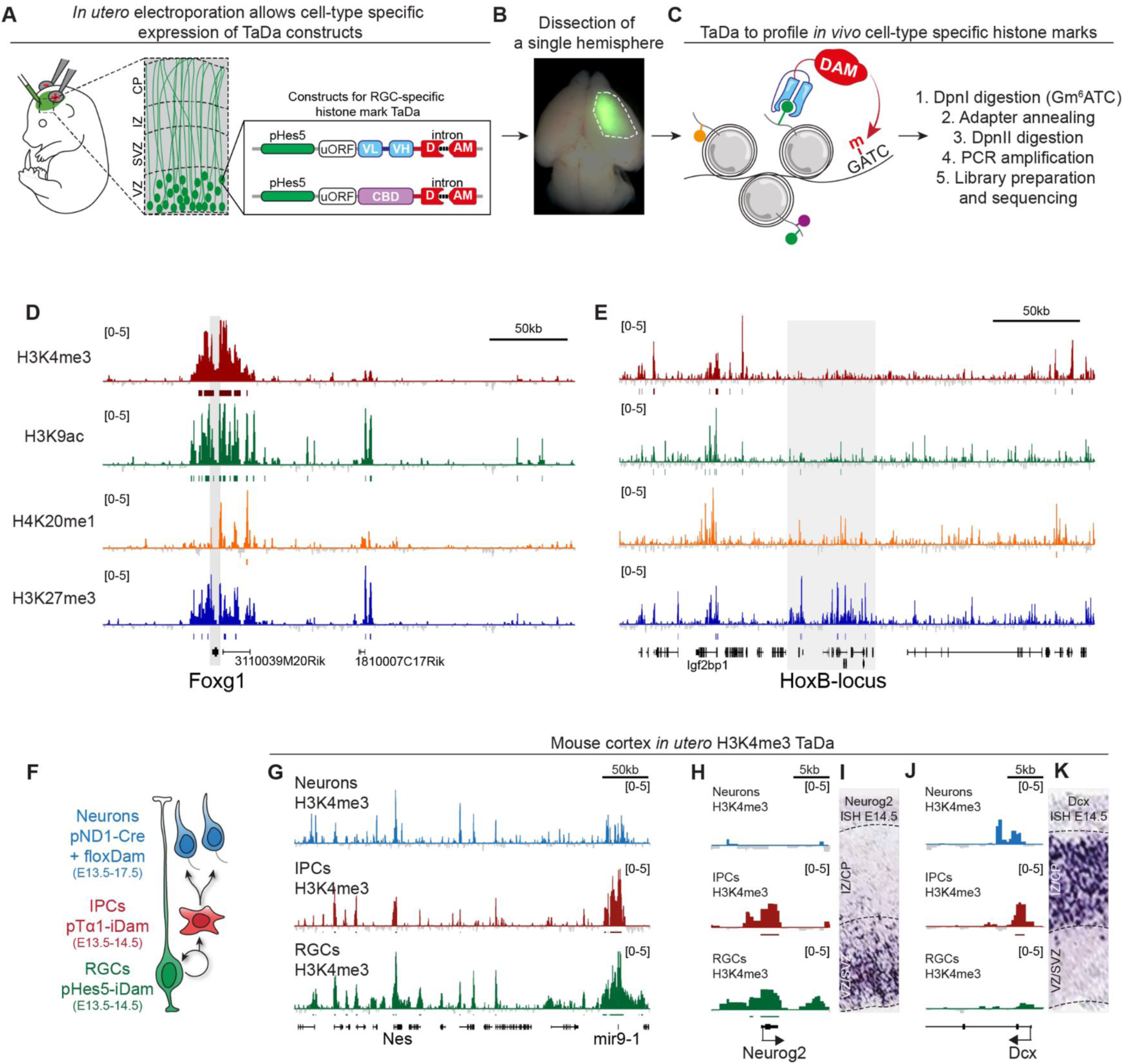
TaDa in mouse RGCs. **A-C.** Schematic of *in utero* TaDa in mouse RGCs. (**D**) TaDa profiles at *FoxG1* in mouse RGCs. (**E**) TaDa profiles at the *HoxB* locus in mouse RGCs. (**F**) Schematic of TaDa during mouse cortical neurogenesis. (**G**) H3K4me3 TaDa profiles at the *Nestin* locus throughout neurogenesis. (**H-I**) H3K4me3 TaDa profiles at the *NeuroG2* locus throughout neurogenesis and the mRNA expression pattern of *NeuroG2* at E14.5. (**J-K**) H3K4me3 TaDa profiles at the *Dcx* locus throughout neurogenesis and the mRNA expression pattern of *Dcx* at E14.5.

H4K20me1 was not studied as part of the ENCODE project (ENCODE Project Consortium., 2012), but H4K20me1 TaDa peaks were strongly correlated with modifications that mark transcribed regions (H3K79me2 and H3K36me3), where H4K20me1 was previously shown to be enriched (**Fig. S2E**). The promoters of genes that are highly expressed in mouse RGCs were correlated with the presence of activating H3K4me3 and H3K9ac histone modifications (**Fig. S2H**). However, in contrast to *Drosophila* NSCs, many of these promoters were also labelled by the repressive mark H3K27me3 (**Fig. S2H**). Although TaDa profiles cells on a population level that could display cell-to-cell heterogeneity in transcription and histone modifications, the widespread overlap of activating and repressive marks suggest bivalency of mouse NSC gene promoters, although analysis of H3K4me3/H3K27me3 at a single cell, single allele level would be necessary to prove this conclusively.

### Cell-type specific histone marks throughout cortical neurogenesis

During neurogenesis, RGCs in the developing mouse cortex differentiate into intermediate progenitors (IPCs), and postmitotic neurons that migrate to the cortical plate (**Fig. 2F**). To profile histone modifications during neurogenesis we used a Hes5 promoter-fragment (pHes5 (Mizutani et al., 2007; Ohtsuka et al., 2006)) to restrict H3K4me3 TaDa to RGCs and the promoter of the Tuba1a gene (pTα1 (Gal et al., 2006; Gloster et al., 1994; Mizutani et al., 2007)) to restrict H3K4me3 TaDa to IPCs (**Fig. 2F**). To target postmitotic neurons, we expressed Cre recombinase from a NeuroD1 promoter (pND1 (Hand et al., 2005)) with a floxDam construct (van den Ameele et al., 2022) (**Fig. 2F)**. The combination gave continuous expression of H3K4me3 TaDa after recombination in young postmitotic neurons (**Fig. 2G**).

The histone modification profiles were cell-type specific, as illustrated by identification of H3K4 trimethylation at the promoter of the progenitor-specific gene Neurog2 in RGCs and IPCs, but not neurons (**Fig. 2H,I**). Similarly, the promoter of the neuronal gene Dcx was marked by H3K4me3 in IPCs and neurons, but not in RGCs (**Fig. 2J,K**). TaDa generated accurate genome-wide profiles of histone modifications from small numbers of cells, including transient progenitor populations like IPCs, without dissociating them from the complex cellular environment of the developing cerebral cortex.

### Profiling histone modifications during *Xenopus* embryogenesis

Early development of the *Xenopus* embryo is a powerful system to study cellular differentiation and reprogramming (Gurdon, 1960) and the role of epigenetic modifications (Hörmanseder et al., 2017). ChIP experiments from early-stage embryos require large amounts of starting material and have proven difficult due to the yolk and protein-rich cytoplasm. We generated constructs for *in vitro* transcription of H3K4me3 TaDa mRNA and injected these into the cytoplasm of fertilized eggs. DamID-seq was performed on single gastrulae at stage 11, and on single dissected dorsal (ecto- and mesoderm) or ventral (endoderm) halves of embryos at neurula stage 21 (**Fig. 3A**). Despite having single embryos as starting material, the H3K4me3 profiles were highly sensitive and specific as demonstrated by selective Dam-methylation on the promoters of genes known to be differentially expressed in mesoderm (*Meox2*; **Fig. 3B,C**) or endoderm (*Sox17* locus; **Fig. 3D**). In comparison with ChIP-seq, requiring 50 gastrulae or the tissue from 75 stage 21 embryos (Hörmanseder et al., 2017), TaDa enabled us to profile H3K4me3 genome-wide in a single embryo.

**Figure 3.**
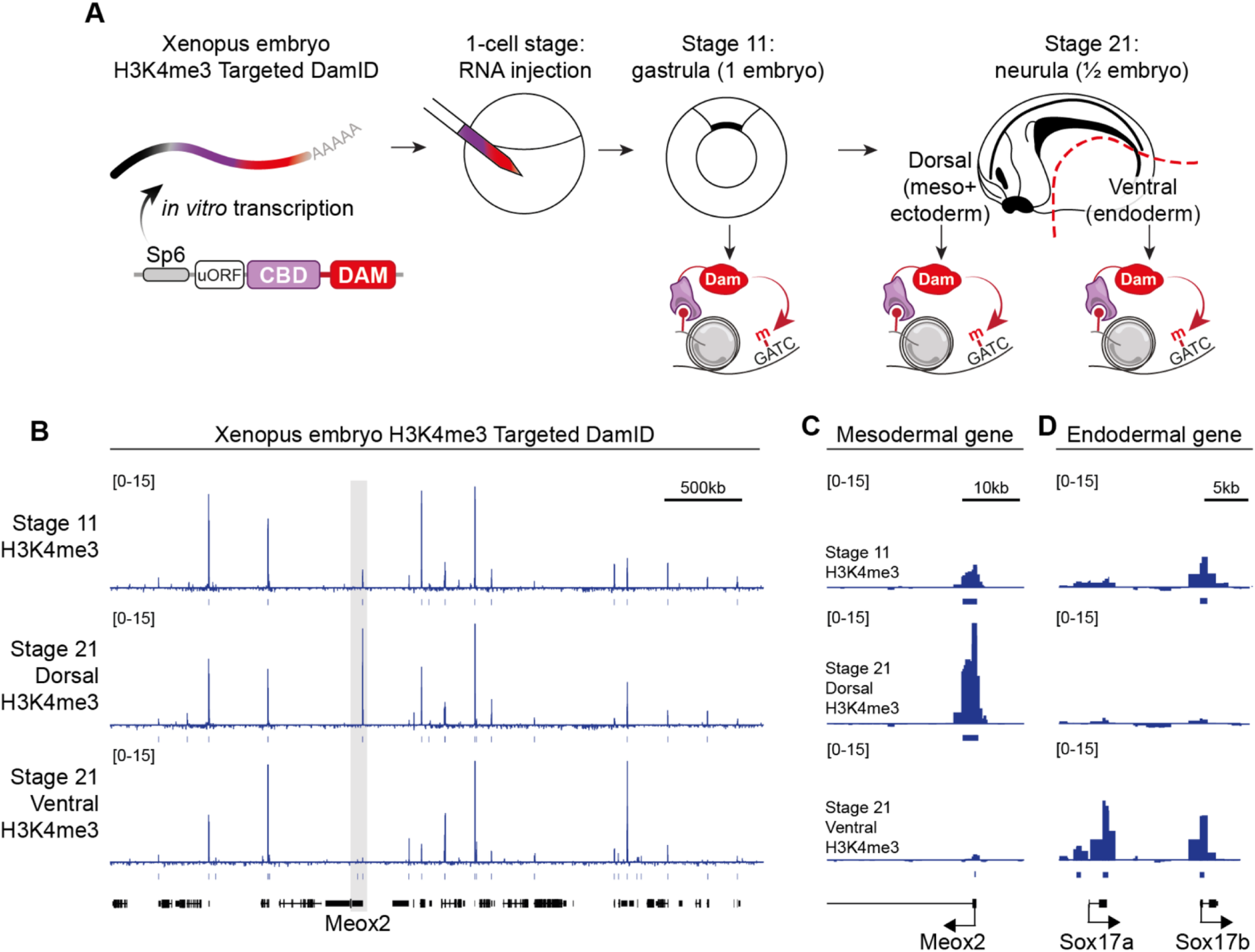
Cell-type specific histone mark TaDa in *Xenopus* embryos. (**A**) Schematic of H3K4me3 TaDa during Xenopus embryogenesis after injection of *in vitro* transcribed mRNA in 1-cell stage embryos. (**B-D**) H3K4me3 TaDa profiles at the indicated developmental stages on the *Meox2* and *Sox17* loci.

## Discussion

Here we show that TaDa can be used to profile posttranslational modifications on histones in a cell type specific manner. Recombinant binders are increasingly used in research (Harmansa and Affolter, 2018), and chromatin readers have been applied previously to optimise ChIP-seq protocols (Hattori et al., 2013; Kungulovski et al., 2014; Kungulovski et al., 2016), or in combination with proximity biotinylation to detect proteins associated with specific histone marks (Villaseñor et al., 2020). Chromatin readers have also been used in combination with DamID to allow the profiling of select histone modifications when combined with single cell sequencing technologies (Rang et al., 2022). Combining the versatility of chromatin readers with the strength of TaDa opens new avenues for studying cell-type specific regulation of chromatin without the necessity for cell dissociation, nuclei extraction or crosslinking. Our results further demonstrate the potential for TaDa to profile any antigen for which a recombinant binder or single chain antibody can be developed, as we recently demonstrated with nanobodies against GFP (NanoDam; Tang et al., 2022). Most of the chromatin readers we tested generated good signals and their distribution across the genome correlated with previously published results, however, this was not the case for all of those tested (e.g. H3K9me3 mintbody) (Hattori et al., 2013). The further development of novel chromatin readers, in particular the optimization of mintbodies for intracellular expression holds great promise in this respect (Chirichella et al., 2017; Harmansa and Affolter, 2018; Sato et al., 2013; Sato et al., 2016).

### Materials and Methods Plasmids

pHes5, a 764bp fragment of the mouse Hes5-promoter including the 5’UTR (Ohtsuka et al., 2006), pTa1, a 1097bp fragment of the mouse Tuba1a promoter (Gal et al., 2006; Gloster et al., 1994; Mizutani et al., 2007), pNeuroD1-Cre and pCAG-Venus were described before (van den Ameele et al., 2022).

#### Dam-fusion proteins

The H3K4me3 and H3K27me3 chromatin binding proteins (TAF3 PHD amino acids 856-929 (Kungulovski et al., 2016), the DNMT3A PWWP domain and CBX7 Chromo Domain amino acids 7-61 (Kungulovski et al., 2014)) were amplified by RT-PCR from cDNA of human ESC-derived NSCs. TAF3 PHD was cloned C-terminal, and CBX7 CD N-terminal of Dam. The single chain antibodies against H3K9ac (19e5; gift of Y. Sato (Sato et al., 2013)), H3K9me3 (JH9-6 from clone 309M3-A; gift of S. Koide (Hattori et al., 2013)) and H4K20me1 (15f11; gift of Y. Sato (Sato et al., 2016)) were amplified by PCR from plasmids and cloned N-terminal of Dam. A nuclear localization signal (2xSV40-NLS) was inserted between Dam and all mintbodies or CBX7 CD and Dam. When this NLS was omitted from 19e5-Dam or placed N-terminal of 19e5, methylation efficiency in *Drosophila* NSCs was reduced (data not shown).

#### Intron-dam constructs

i1Dam and i2Dam were described previously (Wade et al., 2021;van den Ameele et al., 2022). I1Dam is intron 3 of mouse IghE (Lacy-Hulbert et al., 2001), placed between the 3^rd^ and 4^th^ helix (BamHI site) of the DNA-binding domain of the Dam methylase (Horton et al., 2006). I2Dam is a modified version of the Promega chimeric intron sequence (Bothwell et al., 1981; Promega, 2009; Senapathy et al., 1990). Modifications to the intron and the exon sequence at the splice junctions of i2Dam were made to optimize *in silico* predicted splicing efficiency, remove a weak predicted acceptor site and insert an XmaI site. Splice site predictions were performed with NNSPLICE0.9 (Reese et al., 1997) and Spliceport (Dogan et al., 2007). Both constructs were cloned into pCAG-IRES-GFP (pCIG, gift from P. Vanderhaeghen) to obtain pCAG-mcherry-i1/2Dam. Introducing an intron into the Dam CDS has the additional benefit that it greatly improves cloning efficiency, by preventing toxicity in bacteria.

#### FloxDam construct

floxDam was described previously (van den Ameele et al., 2022) and contains Lox71 and Lox61 sites (Albert et al., 1995) respectively upstream of mCherry and within intron2. The two Lox-sites and intervening sequence are inverted to obtain pCAG-flox2Dam. Cre recombinase activity will result in unidirectional reversion of this cassette, thus reconstituting the i2Dam construct. Placing the Lox-site within the intron has the advantage that splicing will remove the remaining lox-site from the Dam ORF, which otherwise interferes with methylation efficiency (data not shown).

#### Xenopus plasmids

Dam and Dam-TAF3phd were cloned into the EcoRI-NotI sites of pCS2-HA+. mRNA was synthetized *in vitro* using the MEGAscript SP6 Kit (Ambion, AM1330M) following the manufacturer’s instructions.

#### Drosophila plasmids

pUAST-attB-mCherry-i2Dam was cloned by replacing Dam in pUAST-attB-LT3-Ndam (Southall et al., 2013) with i2Dam from pHes5-mCherry-i2Dam. All Dam-fusion proteins in *Drosophila* were fused to Dam, apart from the CBX7 Chromo Domain, which was fused to i2Dam.

Dam-positive bacteria were 10-beta Competent *E. coli (NEB* C3019H). Dam-negative bacteria were *dam^-^/dcm^-^* competent E. coli (NEB C2925H). Plasmids for IUE and transient transfection DamID were prepared from Dam-negative bacteria with Endofree Plasmid Maxi kit (Qiagen 12362).

### Mice and *in utero* electroporation (IUE)

All mouse husbandry and experiments were carried out in a Home Office-designated facility, according to the UK Home Office guidelines upon approval by the local ethics committee (Project Licence PPL70/8727). Experiments were done in wild-type MF1 mice. Timed natural matings were used, where noon of the day of plug-identification was E0.5. IUE was performed as previously described (Dimidschstein et al., 2013; Saito and Nakatsuji, 2001) at E13.5 with 50ms, 40V unipolar pulses (BTX ECM830) using CUY650P5 electrodes (Sonidel). DamID plasmids were injected at 1µg/µl together with pCAG-Venus at 0.25µg/µl. Embryos were harvested after 24 hours (E14.5) for TaDa with pHes5 and pTa1 or after 72 hours (E16.5) for TaDa with pND1-Cre.

### Xenopus husbandry and mRNA injection

Mature *Xenopus laevis* males and females were obtained from Nasco. Work with *Xenopus laevis* is covered under the Home Office Project Licence PPL70/8591. Frog husbandry and all experiments were performed according to the relevant regulatory standards, essentially as previously described (Hörmanseder et al., 2017). For mRNA injection, eggs were *in vitro* fertilized, dejellied using 2% Cystein solution in 0.1xMMR, pH 7.8, washed 3 times with 0.1x MMR and transferred into 0.5x MMR for injections. 2-3 pg of mRNA was used per injection. Embryos were cultured at 23°C and collected at gastrula stage 11 or neurula stage 21. Genomic DNA extraction for DamID library preparation was performed using a Qiagen DNAeasy blood and tissue kit, as per manufacturer instructions.

### Fly husbandry and transgenesis

UAS-mCherry-i2Dam was injected into y,sc,v,nos-phiC31;attP40;+ (Bl#25709) and y,sc,v,nos-phiC31;+;attP2 (Bl#25710) embryos. All other *Drosophila* constructs were only injected into y,sc,v,nos-phiC31;attP40;+ (Bl#25709) embryos. Male transgenic TaDa flies were crossed to w^1118^;Worniu-GAL4 (from (Albertson, 2004));Tub-Gal80^ts^ (active in NSCs) virgins. Flies were reared in cages at 25°C, embryos were collected on food plates for 3 hr and transferred to 29°C for 24 hours before dissection at third instar.

### DamID-seq

For *Drosophila* DamID, between 20 and 50 larval brains were dissected. For DamID on mouse cortex after IUE, embryos were cooled on ice, and the electroporated cortex was identified with a fluorescent binocular microscope (**Fig. 2B**). Meninges were removed and the electroporated region microdissected. For Xenopus DamID, 5 injected embryos were pooled for genomic DNA extraction, and one fifth of the genomic DNA was processed for DamID. All samples were processed for DamID as described previously (Marshall et al., 2016). DamID fragments were prepared for Illumina sequencing according to a modified TruSeq protocol (Marshall et al., 2016). All sequencing was performed as single end 50 bp reads generated by the Gurdon Institute NGS Core using an Illumina HiSeq 1500.

### DamID-seq data processing

Analysis of fastq-files from DamID-experiments was performed with the damidseq pipeline script (Marshall and Brand, 2015) that maps reads to an indexed bowtie2 genome, bins into GATC-fragments according to GATC-sites and normalises reads against a Dam-only control. Preprocessed fastq-files from *Drosophila*, *Xenopus* and mouse were mapped respectively to the mm10 (GRCm38.p6), dm6 (BDGP6) or Xla.v9.1 genome assemblies. Reads were binned into fragments delineated by 5’-GATC-3’ motifs (GATC-bins). Individual replicates (see Fig. S1 for n-numbers) for the Dam-fusion constructs were normalized against separate Dam-only replicates with a modified version of the damidseq_pipeline (Marshall and Brand, 2015) (RPM normalization, 300 bp binsize) and all resulting binding profiles for one Dam-fusion construct were quantile normalized to each other (Marshall and Brand, 2015). The resulting logarithmic profiles in bedgraph format were averaged for all GATC-bins across the genome and subsequently backtransformed (“unlog”). Files were converted to the bigwig file format with bedGraphToBigWig (v4) for visualization with the Integrative Genomics Viewer (IGV) (v2.4.19).

### Peak calling and analysis

Macs2 (v2.1.2) (Zhang et al., 2008) was used to call broad peaks for every dam-fusion/dam-only pair on the set of *.bam-files generated by the damidseq_pipeline, using Dam-only as control. Peaks were filtered stringently for FDR<10^-2^ (mouse) or FDR<10^-5^ (*Drosophila* and *Xenopus*) and were only considered if present in all pairwise comparisons for a particular experimental condition. Peaks were merged when they were within 1kb from each other with the merge-function from bedtools (v2.26.0). A similar approach was used for the published ChIP-seq datasets, without control. Enrichment of publicly available histone modification datasets from ENCODE or modENCODE for mouse peak coordinates was determined with i-cisTarget (Imrichová et al., 2015). Overlapping features were detected by uploading peak coordinates to the i-cisTarget online platform (Gene annotation: RefSeq r70; Database: v5.0) after conversion of mouse mm10 peak coordinates to mm9 with the online UCSC liftOver tool. Normalized Enrichment Scores were plotted as bar graphs in Microsoft Excel.

Associating peaks with the defined genomic features was performed using ChIPseeker (v3.10), with plotAnnoBar function and promoter defined as 2kb upstream of the transcription start site.

### Genome wide correlation

Correlation of signal intensity between TaDa conditions and with ChIP-seq datasets from mouse E14.5 forebrain (ENCODE) across the genome was done with the multiBigwigSummary and plotCorrelation functions from the deepTools suite (v3.1.3).

Default bin size of 10kb was used, which allows smoothing of the signal and accounts for variability in peak centre location between TaDa and ChIP-seq. For genome-wide correlation between replicates a bin size of 1kb (-bs 1000) was used. Spearman’s correlation coefficient was calculated to account for large differences in signal intensity between TaDa and ChIP-seq.

### Gene coordinates and gene expression data

*Drosophila* NSC gene expression came from (Yang et al., 2016). For genes expressed in NSCs, all genes with TPM>10 in all three replicates from sorted antennal lobe NSCs were included; for genes not expressed in NSCs, all genes with 0≤TPM<1 in all three replicates were included. Mouse RGC gene expression came from (Florio et al., 2015). The top or bottom 1500 genes from the aRG bulk RNA seq dataset (average of 4 replicates) were used for further analysis. Mouse gene coordinates of protein-coding transcripts defined by ensembl were downloaded from biomaRt (v2.38.0). *Drosophila* gene coordinates of “canonical genes” were obtained from the UCSC Table Browser (Karolchik et al., 2004).

### Data visualization

Genome browser views were generated using the Integrative Genomics Viewer IGV (v2.4.19) (Robinson et al., 2011) with the midline for TaDa ratio tracks set at 1 and for ChIP-seq set at 0. Peak or gene coordinates were saved in .bed format and supplied as features to Seqplots (v1.12.1) (Stempor and Ahringer, 2016); quantile normalized, averaged and backtransformed TaDa profiles were provided in bigwig format. plotAverage and plotHeatmap were used to visualize and average the binding intensities across all supplied coordinates with bin-size of 50bp. Clustering of histone mark signal over gene bodies (including 5kb up- and downstream) was done with the built-in clustering tool of SeqPlots (v1.12.1) (Stempor and Ahringer, 2016), using k-means clustering into 10 separate clusters. Figures were assembled in Adobe Illustrator.

### Previously published datasets

In situ hybridization images were from the Eurexpress database (Diez-Roux et al., 2011). Histone mark ChIP-seq datasets from mouse E14.5 forebrain (ENCFF002AME for H3K4me3; ENCFF002ANR for H3K9ac; ENCFF002DQB for H3K9me3; ENCFF002AFY for H3K27me3) was downloaded respectively from the ENCODE (Bernstein et al., 2012) portal as fastq-files and remapped to the mm10 genome assemblies using the just_align function of the damidseq pipeline (Marshall and Brand, 2015). *Drosophila* NSC RNA PolII binding data came from GSE77860 (Marshall et al., 2017). Mouse RGC gene expression came from GSE65000 (Florio et al., 2015).

## Data availability

All sequencing data is available from GEO.

## Acknowledgements

We thank Y. Sato, S. Koide, P. Vanderhaeghen and W. Gruhn for plasmids and ESC derived NSC cDNA; K. Harnish (Gurdon Institute NGS Core facility) for help with sequencing; the Gurdon Institute animal facility; C. Bradshaw for advice on bioinformatic analysis, and members of the Brand lab for comments on the manuscript. This work was funded by Wellcome Trust Senior Investigator Award (103792) and Royal Society Darwin Trust Research Professorship to AHB. JvdA was supported by a Wellcome Trust Postdoctoral Training Fellowship for Clinicians (105839). EH was supported by a Chromatin Dynamics (CRC1064) and Project Grant (HO 6864/2-1) from the German Research Foundation, Marie Sklodowska-Curie Postdoctoral Fellowship and EMBO Long-Term Fellowship for Postdoctoral Research. JBG and EH were funded by the Medical Research Council (MR/P000479/1). AHB acknowledges core funding to the Gurdon Institute from the Wellcome Trust (092096) and CRUK (C6946/A14492).

## Author contributions

JvdA, SWC, EH and AHB designed experiments. JvdA, SWC, MT, EH and RY performed experiments. JvdA, APAD, OLB, RK and SWC performed bioinformatic analysis. JvdA, SWC, OLB, APAD and AHB wrote the paper, with input from all authors. JBG, EH and AHB supervised the project.

## Supplementary figure legends

**Supplementary Fig. 1.**
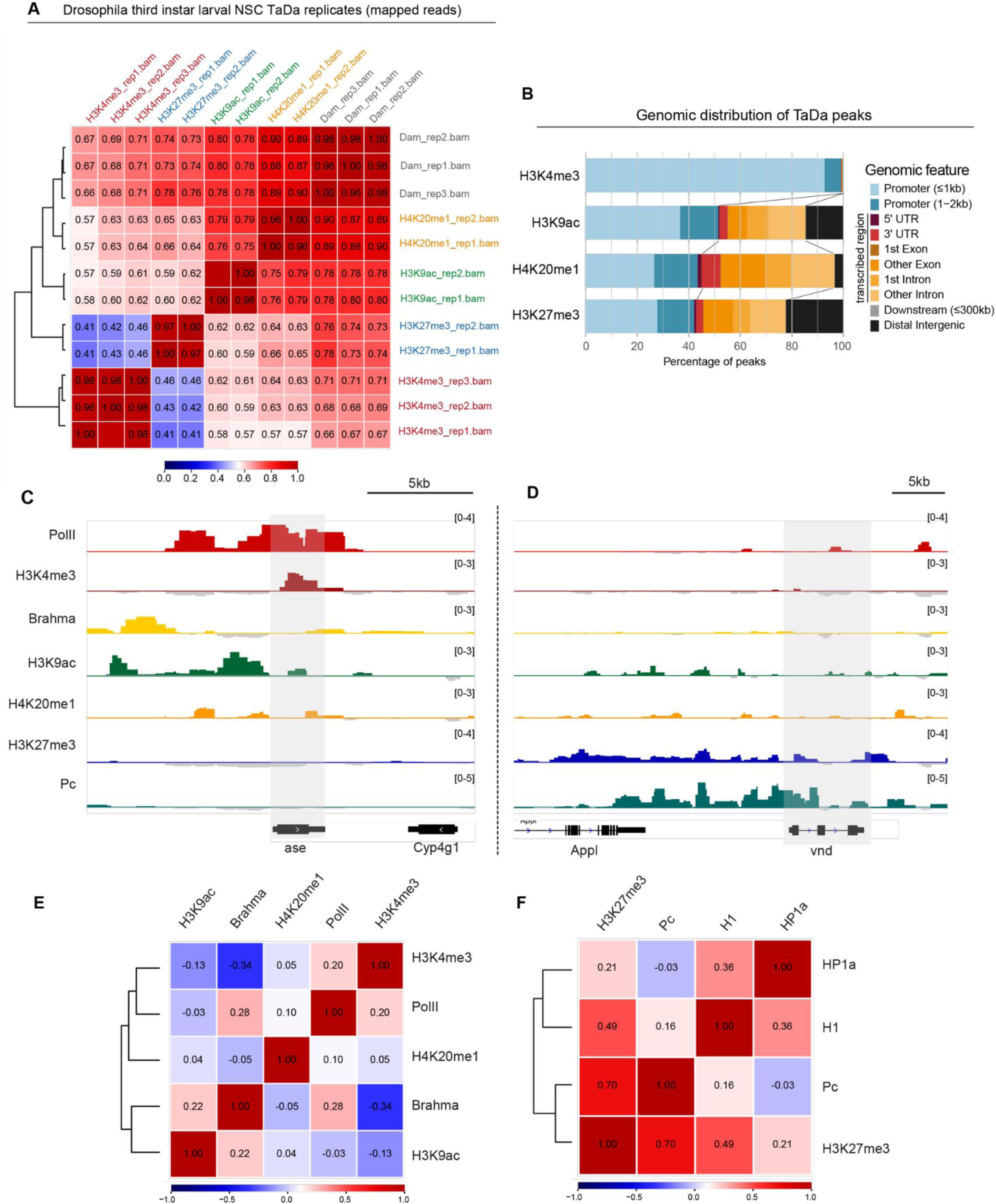
TaDa of histone modifications in Drosophila third instar larval NSCs. (**A**) Pearson correlation coefficients between all TaDa replicates from Drosophila third instar larval NSCs and Dam-only conditions. (**B**) Genomic feature distribution of TaDa profiles in Drosophila third instar larval NSCs. (**C**) TaDa profiles in Drosophila third instar larval NSCs at the *asense* locus (shaded). (**D**) TaDa profiles in Drosophila third instar larval NSCs at the *vnd* locus (shaded). (**E-F**) Pearson correlation coefficients between TaDa using chromatin readers and previously profiled chromatin binding proteins (Marshall & Brand., 2017).

**Supplementary figure 2.**
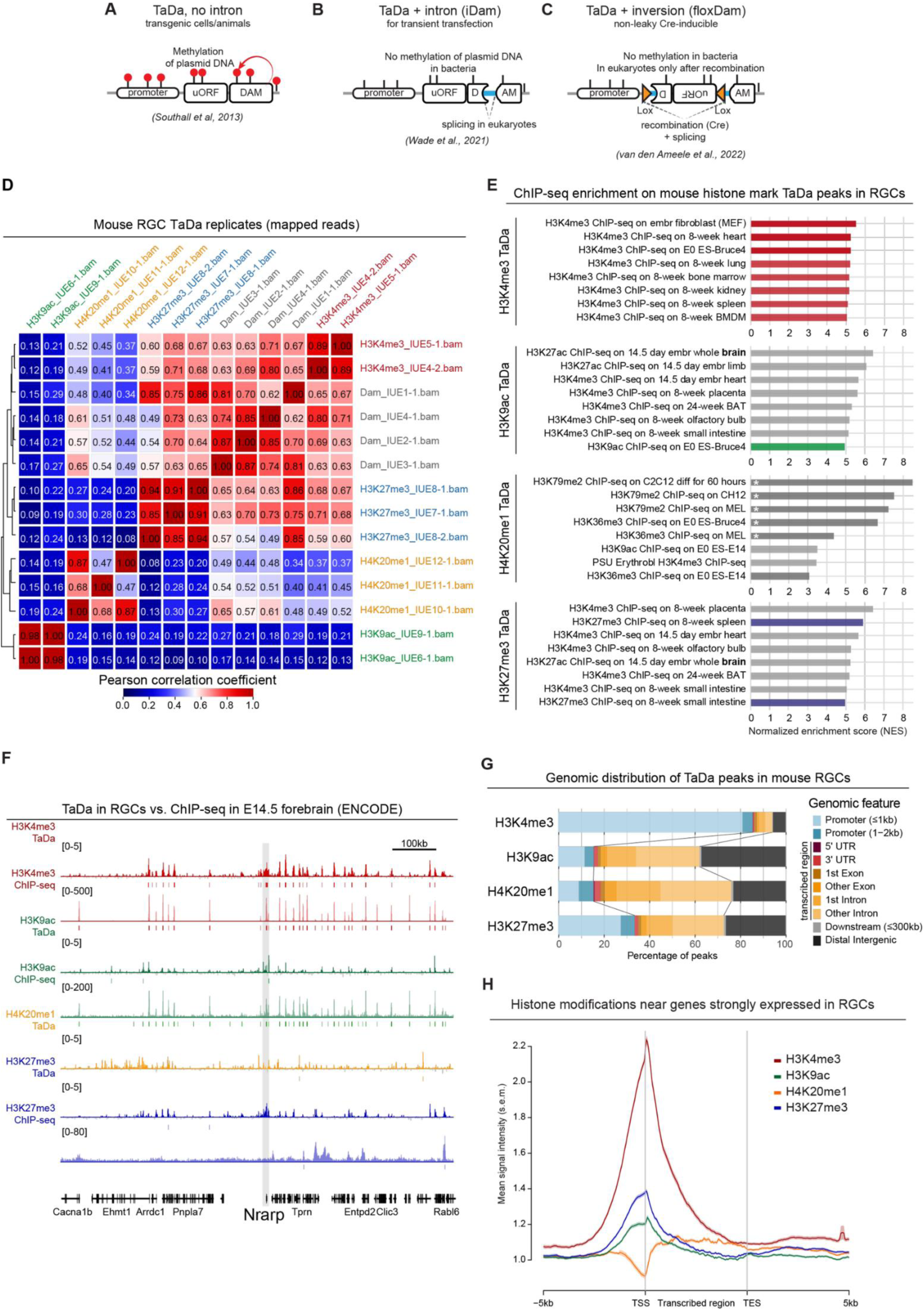
TaDa of histone modifications in mouse RGCs. (**A-C**) Schematic overview of conventional TaDa constructs and those compatible with transient transfection, by introducing an intron or an inversion flanked by lox-sites for recombination. (**D**) Pearson correlation coefficients between all mouse RGC TaDa replicates and the Dam-only conditions. (**E**) Correlations of TaDa for histone modifications in mouse RGCs to ChIP-seq data from the ENCODE consortium database. (**F**) TaDa (top) and ChIP-seq (ENCODE) profiles for histone modifications near the *Nrarp* genomic locus (shaded). (**G**) Genomic feature distribution of histone mark TaDa profiles in mouse RGCs. (**H**) Average signal intensity (±s.e.m.) of TaDa signal at genes (TSS to TES ±5kb) expressed in mouse RGCs.

